# Plant regulator of flower bud differentiation in *in vitro* plants of *Cymbidium tortisepalum* var. *longibracteatum* with TDZ as the key initiator

**DOI:** 10.64898/2026.07.29.741633

**Authors:** Cheng Qiao Wu, Tian Yuan Zhao, Feng Ying Cao, Hao Li, Jie Li

## Abstract

**Background:** *Cymbidium tortisepalum* var. *longibracteatum* is a Class II nationally protected endangered plant in China with significant economic value.

**Aim:** This study aimed to establish an efficient in vitro flowering system and identify key hormonal factors regulating flower bud differentiation.

**Methods:** Orthogonal experiments and hormone treatments were performed to evaluate the effects of TDZ, 6-BA, NAA, IBA, PP_333_, ABA, and GA_3_ on flower bud induction in tissue-cultured plantlets.

**Results:** TDZ was identified as the key initiator of flower bud differentiation, as no flower buds were induced in TDZ-free treatments. The optimal hormone combination was 1.0 mg^·^L^−1^ 6-BA + 0.4 mg^·^L^−1^ TDZ + 0.8 mg^·^L^−1^ NAA + 1.6 mg^·^L^−1^ IBA, achieving a flower bud induction rate of 23.33% and a normal flower bud rate of 20.00%. PP_333_ inhibited flower bud differentiation but reduced malformation; at 0.2 mg^·^L^−1^ PP_333_, the normal flower bud rate reached 20.00%. GA_3_ pretreatment resulted in a flower bud induction rate of 12.04%, whereas ABA pretreatment showed no significant promoting effect on flower bud induction.

**Conclusions:** This study provides an efficient in vitro flowering system and key hormonal parameters for shortening the breeding cycle and elucidating orchid flowering mechanisms.

## Introduction

The family Orchidaceae is one of the largest families of monocotyledonous plants, comprising more than 800 genera and over 30,000 species (Chase et al., 2015). *C. tortisepalum* var. *longibracteatum* is a variety of *Cymbidium tortisepalum* and is recognized as one of the seven major groups of Chinese orchids cultivated in mainland China (Chen and Liu, 2003). It is characterized by upright sword-shaped leaves, evergreen foliage, elegant flowers, and a delicate, long-lasting fragrance, which contribute to its considerable ornamental and economic value (Wang et al., 2025). As an important germplasm resource, wild populations of *C. tortisepalum* var. *longibracteatum* have been included in the International Union for Conservation of Nature (IUCN) Red List of Threatened Species. In commercial production, *C. tortisepalum* var. *longibracteatum* is mainly propagated by division; however, this method has a low multiplication rate, a long propagation cycle, and strong dependence on geographical and seasonal conditions (Zhou et al., 2013). Moreover, the breeding cycle for new *C. tortisepalum* var. *longibracteatum* cultivars is extremely lengthy. Generally, approximately 3 years are required from pollination to seedling establishment, and an additional 7–10 years of cultivation are needed before flowering, which severely limits breeding efficiency. Therefore, accelerating breeding and shortening the breeding cycle are of great significance for cultivar improvement and meeting the increasing demands of the orchid indus try.

In vitro flowering induction is a technique that promotes the transition of tissue-cultured plantlets from vegetative growth to reproductive development by manipulating culture conditions and medium composition (Teixeira da Silva et al., 2014). Previous studies have demonstrated that flowering is coordinately regulated by multiple factors, including environmental cues, nutritional status, and phytohormones, and that the underlying mechanism primarily depends on the synthesis, transport, and distribution of endogenous hormones and assimilates. Therefore, exogenous application of plant growth regulators can be used to break dormancy and induce flowering (Long et al., 2021). At present, studies on in vitro flowering in Orchidaceae have mainly focused on tropical orchids, such as species of *Oncidium* (Kerbauy, 1984), *Phalaenopsis* (Duan and Yazawa, 1995), and *Dendrobium* (Campos an d Kerbauy, 2004; Teixeira da Silva et al., 2014; Lee and Chen, 2014). In contrast, research on in vitro flowering of Chinese orchids remains limited. To date, successful induction of in vitro flowering has only been reported in a few Chinese orchid taxa, including *Cymbidium goeringii* (Zhi et al., 2025), *C. ensifolium* (Wang et al., 1988), *C. kanran* (Zhu et al., 2008), hybrids of *C. goeringii* × *C. hybrida* (Zheng and Pang, 2006), *C. goeringii* × *C*. ‘Hanxiangmei’ (Zeng et al., 2021), and *C. hybridium* ‘Dafeng’ × *C. serratum* var. *goer ingii* ‘Taijishengmei’ (Zhang et al., 2023). Although in vitro flowering has been achieved in species such as *C. goeringii, C. ensifolium*, and *C. kanran*, existing systems generally suffer from low flowering efficiency, high rates of floral abnormalities, and poor reproducibility. More importantly, there are significant differences in how genotypes respond to plant growth regulators, and it is difficult for a single formulation to be universally applicable across species. For *C. tortisepalum* var. *longibracteatum*, an important germplasm resource with both high economic value and conservation significance, no stable and efficient in vitro flowering system has yet been reported. Furthermore, the key hormonal regulators governing flower bud differentiation and their optimal combinations remain unclear.

Plant growth regulators (PGRs) are critical inducers of floral bud initiation and subsequent development in orchids cultured in vitro. Different orchid species exhibit distinct requirements for hormone types and concentrations. This study aims to shorten the juvenile phase of *C. tortisepalum* var. *longibracteatum* and induce normal flowering under in vitro conditions. In this study, in vitro plantlets of *C. tortisepalum* var. *longibracteatum* were used to investigate the effects of different plant growth regulators and their combinations on in vitro flowering induction and to screen for the optimal hormone formulation for inducing in vitro flowering. The results were further compared with the established flowering system reported for *C. goeringii*, a closely related species. This study provides technical support and a theoretical reference for establishing a stable, highly reproducible, and efficient in vitro flowering system for orchid plantlets.

## Materials and Methods

### Plant material

Rhizomes of *C. tortisepalum* var. *longibracteatum* were used as explants to induce in vitro plantlets. The rhizome induction and proliferation medium consisted of MS medium supplemented with 2.0 mg^·^L^−1^ NAA + 0.5 mg^·^L^−1^ 6-BA + 30 g^·^L^−1^ sucrose + 7.6 g^·^L^−1^ agar, pH 5.8–6.0. After 60 days of culture, the rhizomes were transferred to hormone-free 1/2 MS medium and cultured in vitro for 30 days to strengthen the seedlings. Healthy plantlets, 5–6 cm tall and bearing 3–4 leaves, were then selected for subsequent flowering induction experiments.

## Methods

### Effects of different auxins and cytokinins on in vitro flowering of C. tortisepalum var. longibracteatum

In vitro plantlets of *C. tortisepalum* var. *longibracteatum* with similar growth status and bearing three to four leaves were selected and inoculated onto MS media supplemented with different concentrations of 6-BA, TDZ, NAA, and IBA to induce in vitro flowering. The aim was to screen for the optimal combination of plant growth regulators to promote flowering in *C. tortisepalum* var. *longibracteatum* plantlets. The plant growth regulators tested were classified by function: cytokinins included 6-BA and TDZ (TDZ al so possesses auxin-like activity), while auxins included NAA and IBA. The experiment was designed using an L_25_(5^6^) orthogonal array, with four factors—6-BA, TDZ, NAA and IB A—each having five concentration levels: 6-BA (0, 1, 3, 5, 7 mg • L^−1^), TDZ (0, 0.2, 0.4, 0.6, 0.8 mg • L^−1^), NAA (0, 0.2, 0.4, 0.8, 1.6 mg • L^−1^) and IBA (0, 0.2, 0.4, 0.8, 1.6 mg • L^−1^). All media were supplemented with 30 g^·^L^−1^ sucrose and 7.6 g · L^−1^ agar, pH 5.8–6.0. Each treatment consisted of five culture bottles, with two plantlets inoculated per bottle, and each treatment was replicated three times.

### Effects of different concentrations of PP_333_ on in vitro flowering of C. tortisepalum var. longibracteatum

MS medium containing 1.0 mg • L^−1^ 6-BA + 0.4 mg • L^−1^ TDZ + 0.8 mg • L^−1^ NAA + 1.6 mg • L^−1^ IBA + 30 g • L^−1^ sucrose + 7.6 g • L^−1^ agar, with the pH adjusted to 5.8–6.0, was used as the base medium. In vitro plantlets of *C. tortisepalum* var. *longibracteatum* were cultured on this medium supplemented with different concentrations of PP_333_ (0, 0.2, 0.4, 0.6, 0.8, and 1.0 mg · L^−1^) to investigate its effects on in vitro flowering. Each treatment consisted of five culture bottles with two plantlets per bottle, and the experiment was replicated three times.

### Effects of GA_3_ pretreatment on in vitro flowering of C. tortisepalum var. longibracteatum

In vitro plantlets of *C. tortisepalum* var. *longibracteatum* were inoculated onto MS medium supplemented with 30 g · L^−1^ sucrose and 7.6 g · L^−1^ agar, adjusted to pH 5.8–6.0. The medium was supplemented with GA_3_ at different concentrations (0.2, 0.6, and 1.0 mg · L^− 1^), and the plantlets were pretreated for different durations (20, 40, and 60 days). Subsequently, the plantlets were transferred to MS medium supplemented with 1.0 mg • L^−1^ 6-BA + 0.4 mg • L^−1^ TDZ + 0.8 mg • L^−1^ NAA + 1.6 mg • L^−1^ IBA for continuous culture. Each treatment consisted of five culture bottles, with two in vitro plantlets inoculated per bottle, a nd each treatment was replicated three times.

### Effects of ABA pretreatment on in vitro flowering of C. tortisepalum var. longibracteatu m

In vitro plantlets of *C. tortisepalum* var. *longibracteatum* were first pretreated on MS medium supplemented with 30 g · L^−1^ sucrose and 7.6 g · L^−1^ agar, pH 5.8–6.0, with different concentrations of ABA (0.2, 0.6, and 1.0 mg · L^−1^) for different durations (20, 40, and 60 days). Following the pretreatment, the plantlets were transferred to MS medium supplemented with 1.0 mg • L^−1^ 6-BA + 0.4 mg • L^−1^ TDZ + 0.8 mg • L^−1^ NAA + 1.6 mg • L^−1^ IBA for continuous culture. Each treatment consisted of five culture bottles, with two plantlets inoculated per bottle, and each treatment was replicated three times.

### Culture conditions

All cultures were maintained at 26 ± 1 °C under a 12 h · d^−1^ photoperiod with a light intensity of 1,500–2,000 lx.

### Data analysis

After 90 days of culture, the floral bud induction rate and the normal floral bud formation rate were recorded. Flower buds of *C. tortisepalum* var. *longibracteatum* were evaluated by direct morphological observation. Flower buds with intact floral organs and typical horticultural developmental characteristics of *C. tortisepalum* var. *longibracteatum* were considered normally developed flower buds. Data were analyzed using Microsoft Excel 2021 (*Microsoft Corp*., Redmond, WA, USA). Analysis of variance (ANOVA) and multiple comparisons were performed using SPSS 27.0 software (*IBM Corp*., Armonk, NY, USA). Duncan’s multiple-range test was used for mean separation, with α = 0.05. Origin 2024 (*OriginLab Corp*., Northampton, MA, USA) was used to generate the graphs and charts.

Flower bud induction rate (%) = (Number of plantlets with induced floral buds / Total number of inoculated plantlets) × 100

Normal flower bud formation rate (%) = (Number of plantlets with normally developed floral buds / Total number of inoculated plantlets) × 100

## Results

### Effects of different auxins and cytokinins on in vitro flowering of C. tortisepalum var. longibracteatum

As shown in Table 1, no flower buds were induced in the hormone-free control group. In contrast, some treatments supplemented with plant growth regulators showed significantly higher flower bud induction rates than the control group (*P* < 0.05). Al l treatments without TDZ exhibited a flower bud induction rate of 0, indicating that TDZ plays a critical role in the initiation of flower bud differentiation in *C. tortisepalum* var. *longibracteatum*. Treatments containing 0.8 mg · L^−1^ TDZ were conducive to flower bud differentiation but resulted in a high proportion of malformed flowers (Fig. 2*C, D*). Treatment 19 achieved the highest flower bud induction rate of 26.67%, with a normal flower bud rate of merely 10.00%. Treatment 8 exhibited a flower bud induction rate of 23.33% and a normal flower bud rate of 20.00%, which was numerically higher than all other treatments; although the normal flower bud rate showed no statistically significant difference compared with Treatment 4 (13.33%) and Treatment 10 (12.50%), it represented the maximum value observed across the entire experiment. Therefore, Treatment 8 was identified as the optimal formulation, based on both total bud induction and the proportion of normally formed floral buds. The optimal combination of plant growth regulators for in vitro flowering of *C. tortisepalum* var. *longibracteatum* consisted of 1.0 mg · L^−1^ 6-BA + 0.4 mg · L^−1^ TD Z + 0.8 mg · L^−1^ NAA + 1.6 mg · L^−1^ IBA.

**Table 1.**
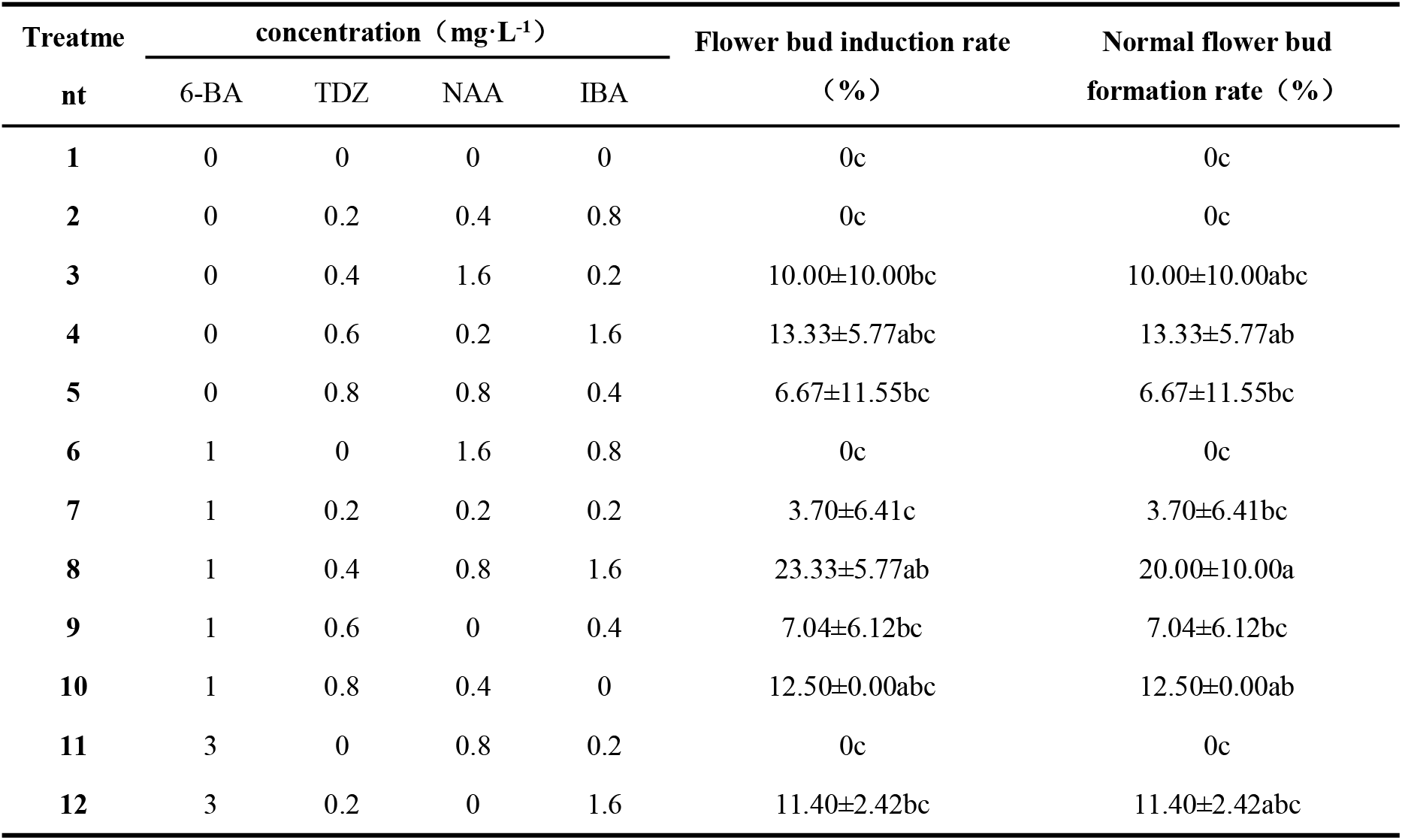
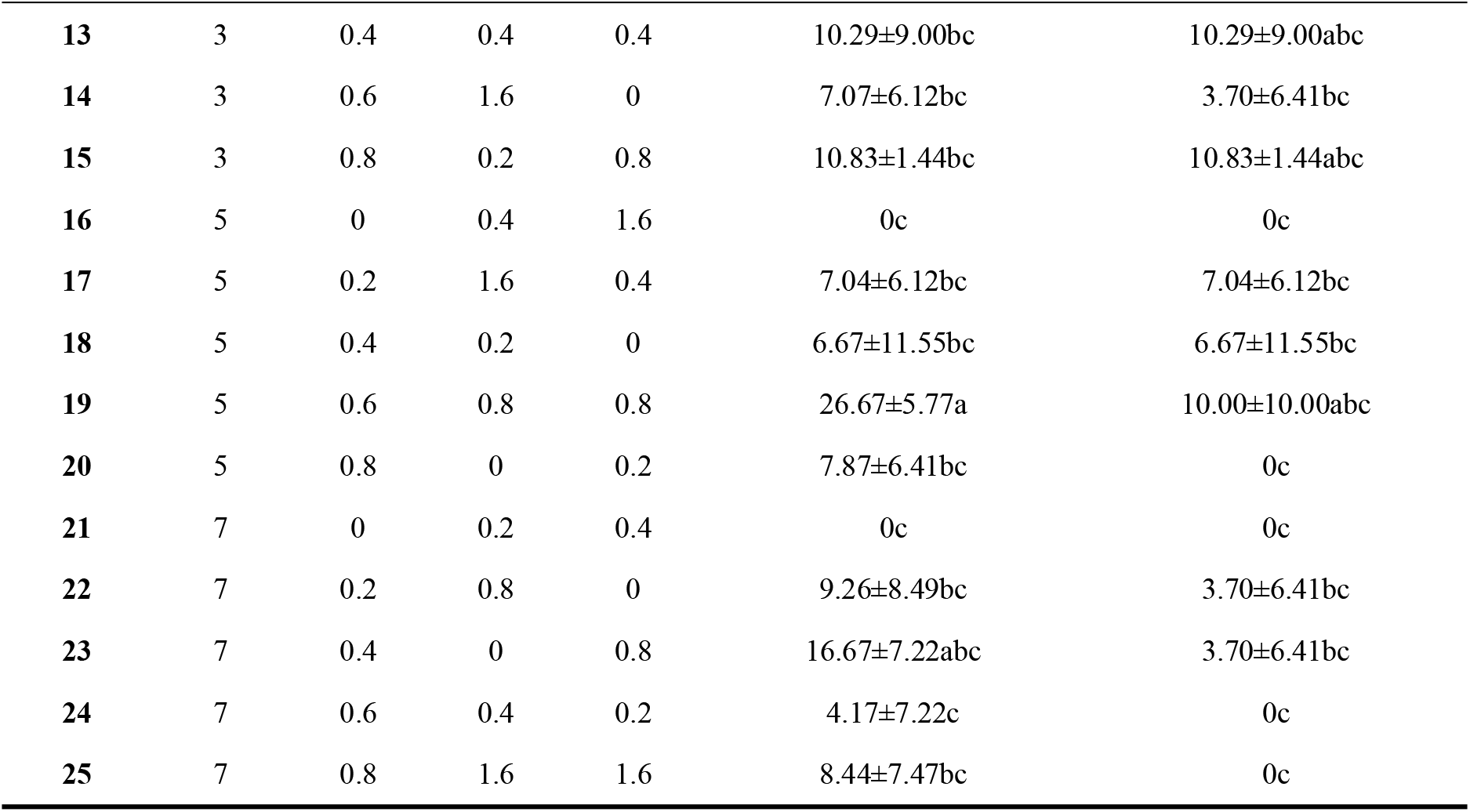
Effects of different auxins and cytokinins on in vitro flowering of *C. tortisepalum* var. *longibrac teatum*. Mean ± SD, n = 3. Different lowercase letters within the same column indicate significant differences between treatments (*P* < 0.05).CK - hormone-free medium.

| Treatment | concentration ($\text{mg}\cdot\text{L}^{-1}$) | | | | Flower bud induction rate (%) | Normal flower bud formation rate (%) |
| --- | --- | --- | --- | --- | --- | --- |
|  | 6-BA | TDZ | NAA | IBA |  |  |
| 1 | 0 | 0 | 0 | 0 | 0c | 0c |
| 2 | 0 | 0.2 | 0.4 | 0.8 | 0c | 0c |
| 3 | 0 | 0.4 | 1.6 | 0.2 | $10.00 \pm 10.00$bc | $10.00 \pm 10.00$abc |
| 4 | 0 | 0.6 | 0.2 | 1.6 | $13.33 \pm 5.77$abc | $13.33 \pm 5.77$ab |
| 5 | 0 | 0.8 | 0.8 | 0.4 | $6.67 \pm 11.55$bc | $6.67 \pm 11.55$bc |
| 6 | 1 | 0 | 1.6 | 0.8 | 0c | 0c |
| 7 | 1 | 0.2 | 0.2 | 0.2 | $3.70 \pm 6.41$c | $3.70 \pm 6.41$bc |
| 8 | 1 | 0.4 | 0.8 | 1.6 | $23.33 \pm 5.77$ab | $20.00 \pm 10.00$a |
| 9 | 1 | 0.6 | 0 | 0.4 | $7.04 \pm 6.12$bc | $7.04 \pm 6.12$bc |
| 10 | 1 | 0.8 | 0.4 | 0 | $12.50 \pm 0.00$abc | $12.50 \pm 0.00$ab |
| 11 | 3 | 0 | 0.8 | 0.2 | 0c | 0c |
| 12 | 3 | 0.2 | 0 | 1.6 | $11.40 \pm 2.42$bc | $11.40 \pm 2.42$abc |
|  | 6-BA | TDZ | NAA | IBA |  |  |
| 13 | 3 | 0.4 | 0.4 | 0.4 | 10.29±9.00bc | 10.29±9.00abc |
| 14 | 3 | 0.6 | 1.6 | 0 | 7.07±6.12bc | 3.70±6.41bc |
| 15 | 3 | 0.8 | 0.2 | 0.8 | 10.83±1.44bc | 10.83±1.44abc |
| 16 | 5 | 0 | 0.4 | 1.6 | 0c | 0c |
| 17 | 5 | 0.2 | 1.6 | 0.4 | 7.04±6.12bc | 7.04±6.12bc |
| 18 | 5 | 0.4 | 0.2 | 0 | 6.67±11.55bc | 6.67±11.55bc |
| 19 | 5 | 0.6 | 0.8 | 0.8 | 26.67±5.77a | 10.00±10.00abc |
| 20 | 5 | 0.8 | 0 | 0.2 | 7.87±6.41bc | 0c |
| 21 | 7 | 0 | 0.2 | 0.4 | 0c | 0c |
| 22 | 7 | 0.2 | 0.8 | 0 | 9.26±8.49bc | 3.70±6.41bc |
| 23 | 7 | 0.4 | 0 | 0.8 | 16.67±7.22abc | 3.70±6.41bc |
| 24 | 7 | 0.6 | 0.4 | 0.2 | 4.17±7.22c | 0c |
| 25 | 7 | 0.8 | 1.6 | 1.6 | 8.44±7.47bc | 0c |

The range analysis (Table 2) results show that the order of influence of the four factors on the flower bud induction rate is TDZ > NAA > IBA > 6-BA, and the optimal combination is 6-BA 5 mg · L^−1^, TDZ 0.4 mg · L^−1^, NAA 0.8 mg · L^−1^, and IBA 1.6 mg · L^−1^. For the normal flower bud rate, the order of influence of the four factors is TDZ > 6-BA > NAA > IBA, and the optimal combination is 6-BA 1 mg · L^−1^, TDZ 0.4 mg · L^−1^, NAA 0. 8 mg · L^−1^, and IBA 1.6 mg · L^−1^. Considering both the flower bud induction rate and the normal flower bud rate, the optimal hormone combination for in vitro flowering of *C. tortisepalum* var. *longibracteatum* is 1 mg · L^−1^ 6-BA + 0.4 mg · L^−1^ TDZ + 0.8 mg · L^−1^ NAA + 1.6 mg · L^−1^ IBA.

**Table 2.**
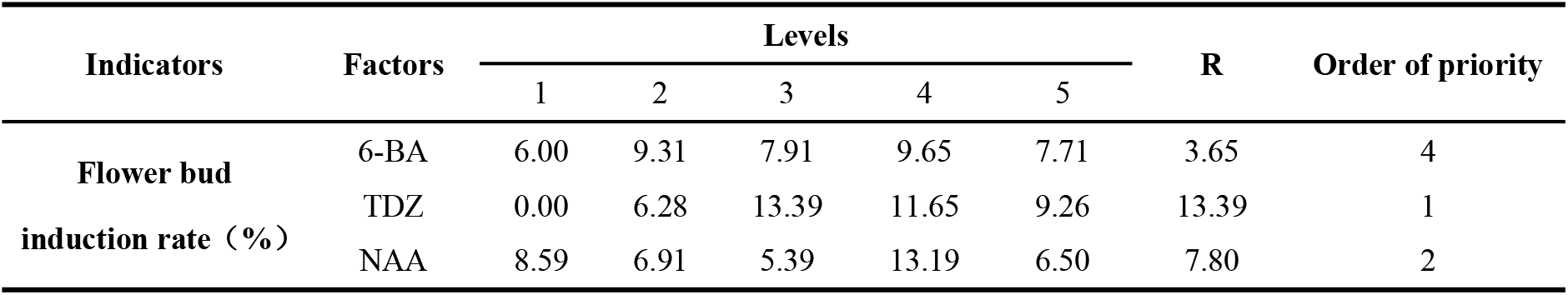
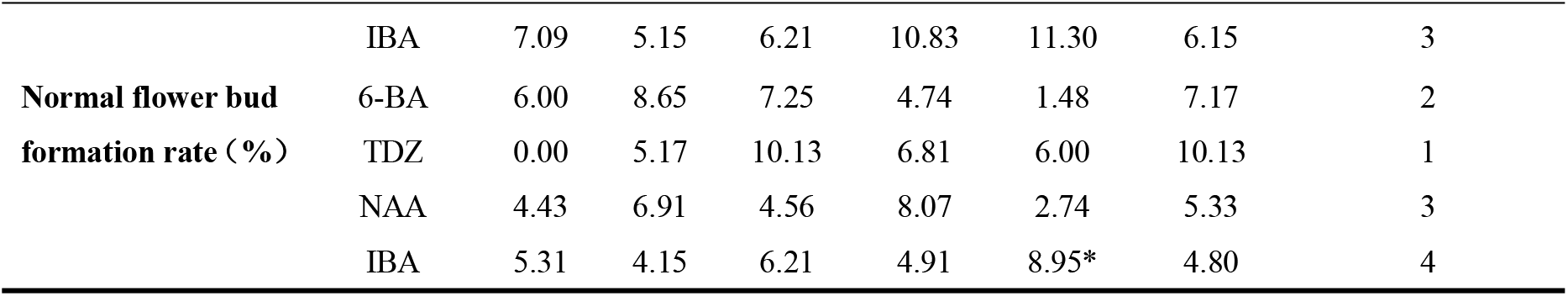
Range analysis of auxin and cytokinin effects on in vitro flowering of *C. tortisepalum* var. *longibracteatum*. The range (R) is used to characterise the extent to which each factor influences the test indicator; a larger R value indicates that the factor has a more significant influence on the test indicator and is a dominant factor, whilst a smaller R value indicates a weaker influence and classifies the factor as a secondary factor.

Analysis of variance(Table 3) revealed that 6-BA had no significant effect on the floral bud induction rate of *C. tortisepalum* var. *longibracteatum*, but significantly affected normal floral bud formation. TDZ had an extremely significant effect on both floral bud induction and normal floral bud formation. NAA significantly influenced floral bud induction, whereas its effect on normal floral bud formation was not significant. Similarly, IBA significantly affected floral bud induction, but had no significant effect on normal floral bud formation.

**Table 3.**
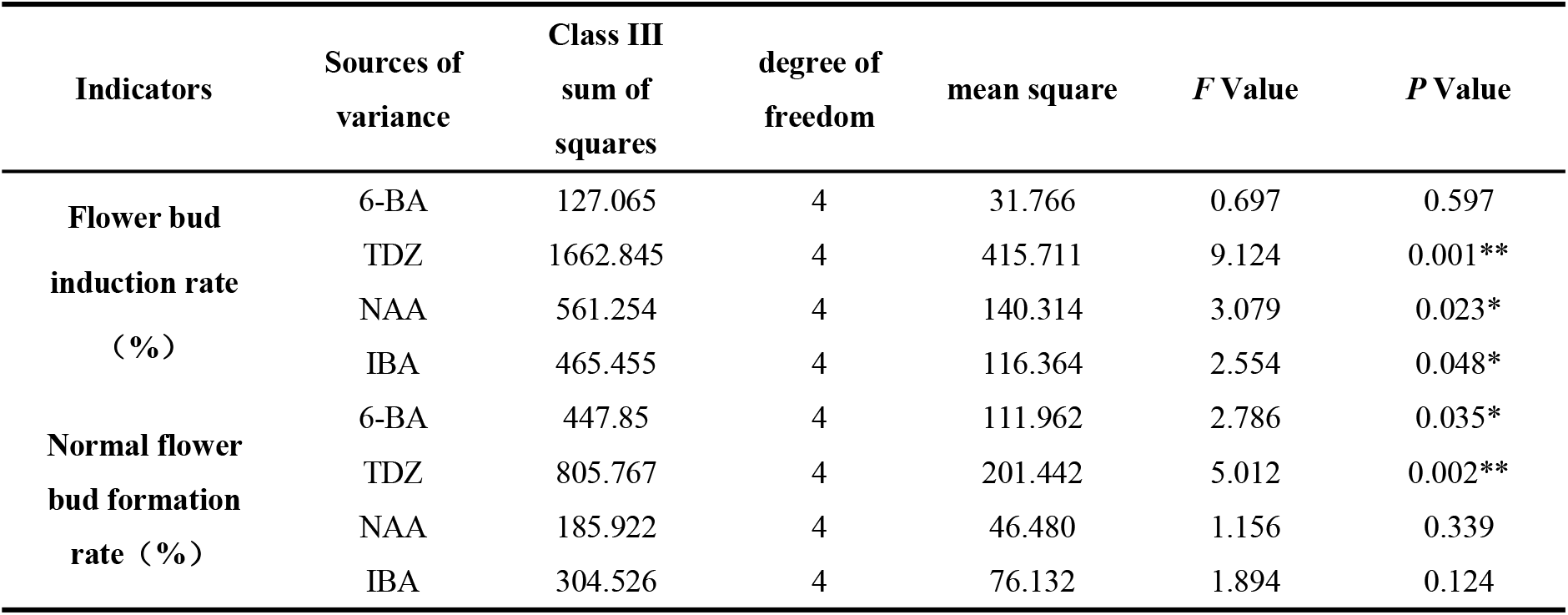
Inter-subject effect test of the effects of auxins and cytokinins on in vitro flowering in *C. tortisepalum* var. *longibracteatum*. (*) indicates a statistically significant effect on flower bud induction rate and normal flowering rate *(P* < 0.05); (**) indicates a highly statistically significant effect on flower bud induction rate and normal flowering rate (*P* < 0.01).

| Indicators | Sources of variance | Class III sum of squares | degree of freedom | mean square | F Value | P Value |
| --- | --- | --- | --- | --- | --- | --- |
| **Flower bud induction rate (%)** | 6-BA | 127.065 | 4 | 31.766 | 0.697 | 0.597 |
|  | TDZ | 1662.845 | 4 | 415.711 | 9.124 | 0.001** |
|  | NAA | 561.254 | 4 | 140.314 | 3.079 | 0.023* |
|  | IBA | 465.455 | 4 | 116.364 | 2.554 | 0.048* |
| **Normal flower bud formation rate (%)** | 6-BA | 447.85 | 4 | 111.962 | 2.786 | 0.035* |
|  | TDZ | 805.767 | 4 | 201.442 | 5.012 | 0.002** |
|  | NAA | 185.922 | 4 | 46.480 | 1.156 | 0.339 |
|  | IBA | 304.526 | 4 | 76.132 | 1.894 | 0.124 |

### Effects of different concentrations of PP_333_ on in vitro flowering of C. tortisepalum var. longibracteatum

MS medium without plant growth regulators was used as the control. The optimal formulation (1.0 mg · L^−1^ 6-BA + 0.4 mg · L^−1^ TDZ + 0.8 mg · L^−1^ NAA + 1.6 m g · L^−1^ IBA) was supplemented with different concentrations of PP_333_. Analysis of variance showed that PP_333_ concentration had an extremely significant effect on the floral bud induction rate and a significant effect on the normal floral bud formation rate (Table 4). As shown in Fig. 1, the floral bud induction rate decreased with increasing PP_333_ concentration. Treatment 1 exhibited the highest floral bud induction rate (23.33%), which was significantly higher than that of all PP_333_-supplemented treatments. The normal floral bud formation rate also decreased with increasing PP_333_ concentration. Both treatments 1 and 2 achieved the highest normal floral bud formation rate (20.00%). Following PP_333_ supplementation, all induced floral buds developed normally, with no obvious malformation observed. Both Treatments 1 and 2 were considered optimal; however, Treatment 1 was selected as the best treatment because it had the highest floral bud induction rate and the lowest probability of malformed flowers. Thus, the optimal formulation was determined to be 1.0 mg · L^−1^ 6-BA + 0.4 mg · L^−1^ TDZ + 0.8 mg · L^−1^ NAA + 1.6 mg · L^−1^ IBA without PP_333_ supplementation.

**Table 4.** Inter-subject effect test of the effects of PP_333_ concentration on in vitro flowering of *C. tortisepalum* var. *longibracteatum*. (*) indicates a statistically significant effect on flower bud induction rate and normal flowering rate *(P* < 0.05); (**) indicates a highly statistically significant effect on flower bud induction rate and normal flowering rate (*P* < 0.01).

| Indicators | Sources of variance | Class III sum of squares | degree of freedom | mean square | F Value | P Value |
| --- | --- | --- | --- | --- | --- | --- |
| Flower bud induction rate (%) | PP<sub>333</sub> | 1495.24 | 6 | 249.206 | 5.815 | 0.003** |
| Normal flower bud formation rate (%) | PP<sub>333</sub> | 1180.95 | 6 | 196.825 | 3.758 | 0.019* |

**Fig. 1.**
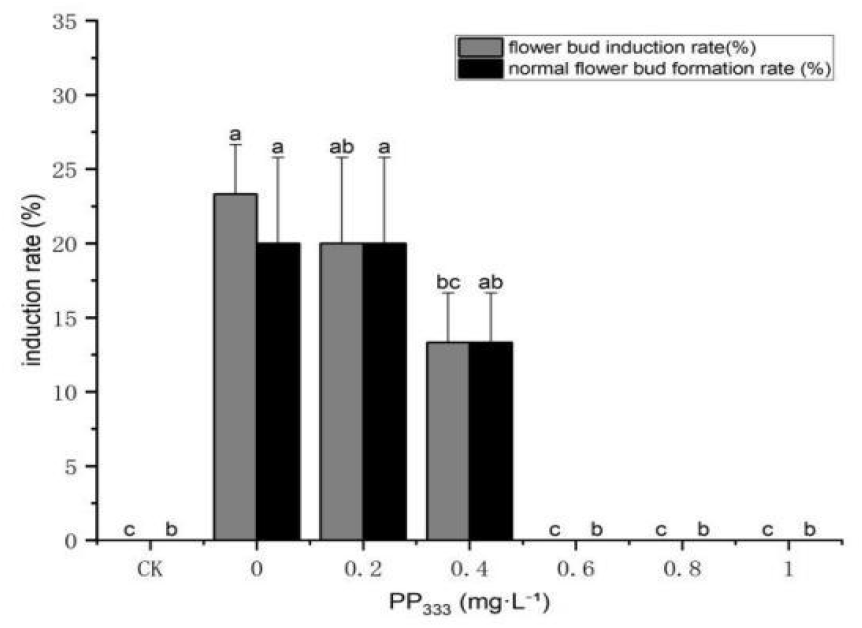
Effects of PP_333_ concentration on in vitro flowering of *C. tortisepalum* var. *longibracteatum*. Mean ± SE, n = 3. Different lowercase letters within the same column indicate significant differences between treatments (*P* < 0.05).CK - hormone-free medium.

**Fig. 2.**
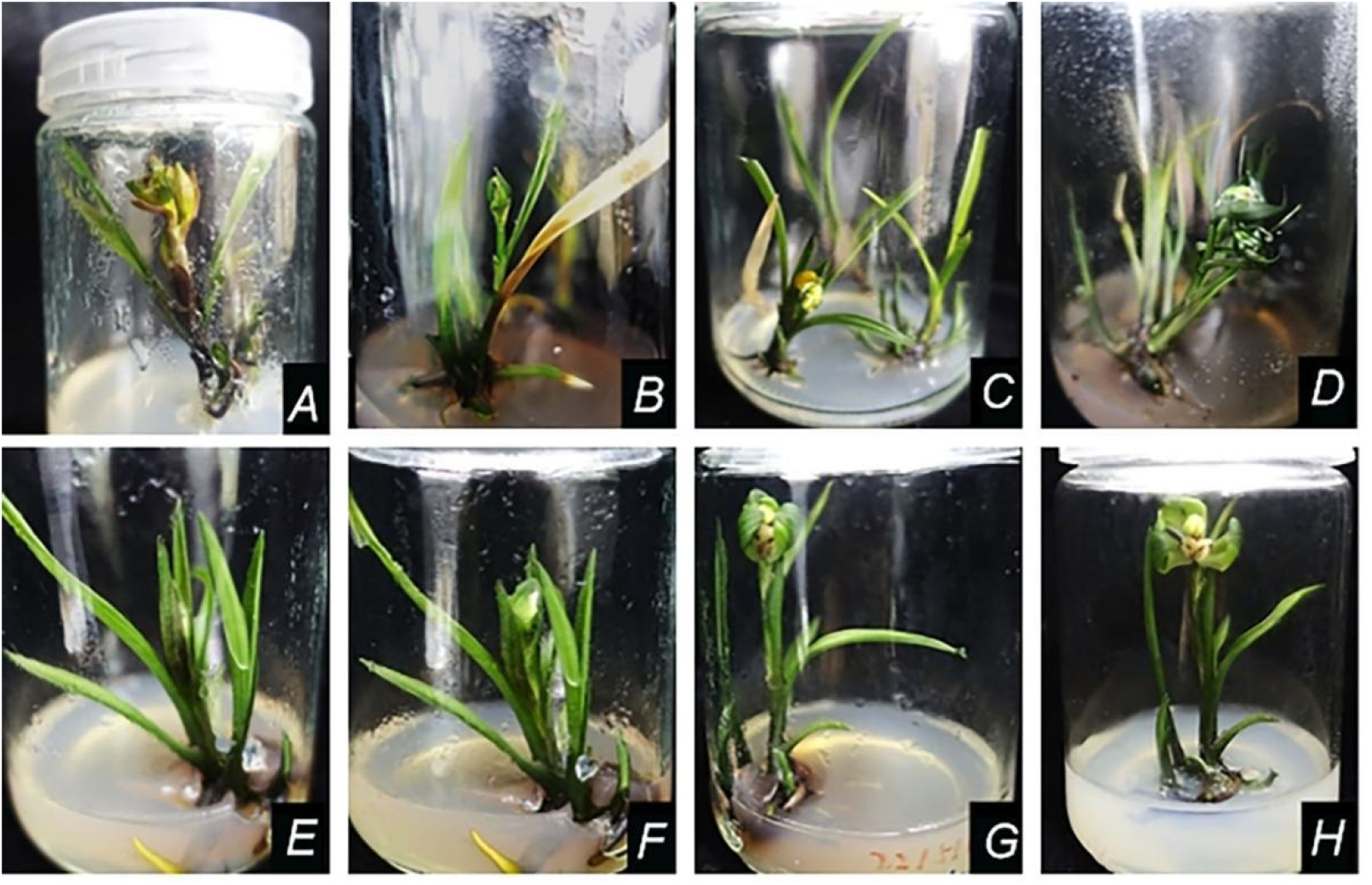
Floral bud induction of *C. tortisepalum* var. *longibracteatum* in vitro induced by plant growth regulators: *A*-*B*: normal flower buds(1.0 mg · L^−1^ 6-BA + 0.4 mg · L^−1^ TDZ + 0.8 mg · L^−1^ NAA + 1.6 mg · L^−1^IBA); *C*: malformed flower buds (5.0 mg · L^−1^ 6-BA + 0.8 mg · L^−1^ TDZ + 0.2 mg · L^−1^ IBA); *D*: semi-open malformed flowers (5.0 mg · L^−1^ 6-BA + 0.8 mg · L^−1^ TDZ + 0.2 mg · L^−1^ IBA); *E*-*H*: the flowering process of the in vitro flower (1.0 mg · L^−1^ 6-BA + 0.4 mg · L^−1^ TDZ + 0.8 mg · L^−1^ NAA + 1.6 mg · L^−1^ IBA).

### Effects of GA_3_ pretreatment on in vitro flowering of C. tortisepalum var. longibracteatu m

Using hormone-free MS medium as the control, different concentrations of GA_3_ (0.2, 0. 6, and 1.0 mg · L^−1^) were added to the MS medium containing the optimal plant growth regulator combination (1.0 mg · L^−1^ 6-BA + 0.4 mg · L^−1^ TDZ + 0.8 mg · L^−1^ NAA + 1.6 mg · L ^−1^ IBA), and the plantlets were pretreated for different durations (20, 40, and 60 days). The results are presented in Table 5. Treatment 6 showed the highest floral bud induction rate and normal floral bud formation rate, both at 12.04%. These values were higher than those of the other treatments. No malformed flowers were observed, indicating good quality of floral bud development. These results suggest that pretreatment with 0.6 mg · L^−1^ GA_3_ for 60 days can effectively induce in vitro flowering in *C. tortisepalum* var. *longibracteatum*.

**Table 5.**
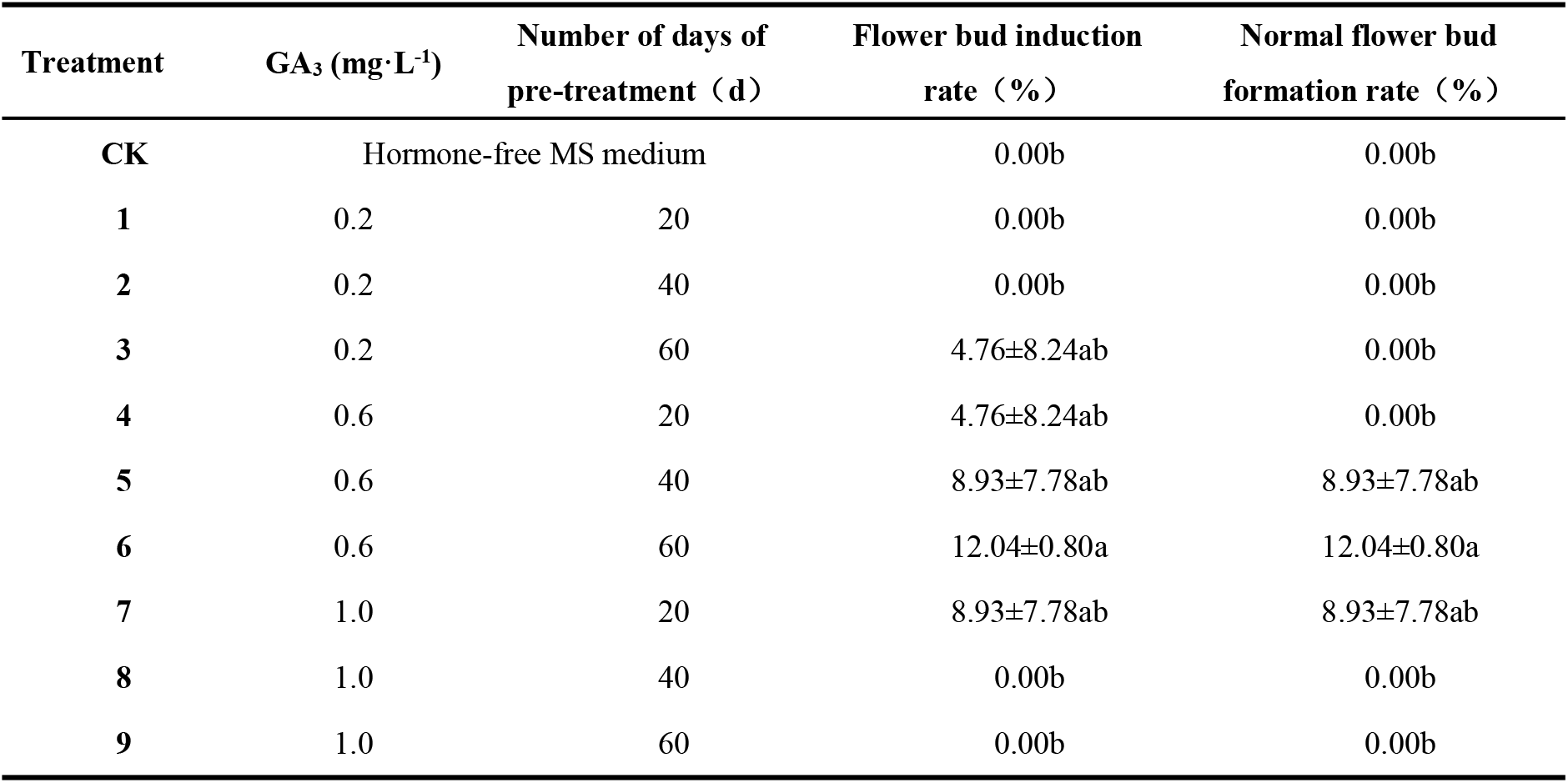
The effect of GA_3_ pretreatment on the in vitro flowering of *C. tortisepalum* var. *longibracteatum*. Mean ± SD, n = 3. Different lowercase letters within the same column indicate significant differences between treatments (*P* < 0.05). CK - hormone-free medium.

The results of the analysis of variance are presented in Table 6. The results indicated statistically significant differences in floral bud induction rate across GA_3_ concentrations, and GA_3_ concentration significantly affected the floral bud induction rate of *C. tortisepalum* var. *longibracteatum*. Moreover, GA_3_ concentration had a highly significant effect on the normal floral bud formation rate, suggesting a more pronounced regulatory role in subsequent floral bud development. In contrast, pretreatment duration had no significant effect on either rate in *C. tortisepalum* var. *longibracteatum*.

**Table 6.** Inter-subject effect test of the effects of GA_3_ pretreatment on the in vitro flowering of *C. tortisepalum* var. *longibracteatum*. (*) indicates a statistically significant effect on flower bud induction rate and normal flowering rate *(P* < 0.05); (**) indicates a highly statistically significant effect on flower bud induction rate and normal flowering rate (*P* < 0.01).

| Indicators | Sources of variance | Class III sum of squares | degree of freedom | mean square | F Value | P Value |
| --- | --- | --- | --- | --- | --- | --- |
| Flower bud induction rate (%) | A: GA<sub>3</sub> | 246.317 | 2 | 123.158 | 3.845 | 0.036* |
|  | B: Number of days of pre-treatment(d) | 31.422 | 2 | 15.711 | 0.491 | 0.618 |
| Normal flower bud formation rate (%) | A: GA<sub>3</sub> | 221.343 | 2 | 110.671 | 9.090 | 0.002** |
|  | B: Number of days of pre-treatment(d) | 6.448 | 2 | 3.224 | 0.265 | 0.770 |

### Effects of ABA pretreatment on in vitro flowering of C. tortisepalum var. longibracteatu m

Using hormone-free MS medium as the control, ABA was added to the MS medium with the optimal formulation (1 mg · L^−1^ 6-BA + 0.4 mg · L^−1^ TDZ + 0.8 mg · L^−1^ NAA + 1. 6 mg · L^−1^ IBA) at 0.2, 0.6, and 1.0 mg · L^−1^, and pre-treatment was carried out for 20, 40, and 60 days. The statistical results are shown in Supplementary Table S1. Among all treatment groups, only Treatment 6 induced flower buds, with an induction rate of 4.55%. However, the induced floral buds failed to develop into fully opened flowers during the later stages of culture.

## Discussion

### Effects of different auxins and cytokinins on in vitro flowering of C. tortisepalum var. longibracteatum

Cytokinins regulate cell division and organ formation and are essential for in vitro flowering in plants (Velmurugan et al., 2010). Cytokinins are synthesized by root tip cells and transported via the xylem to the aboveground parts, where they regulate the transition from vegetative to reproductive growth (Kumari et al., 2023). 6-BA, a commonly used cytokinin, effectively promotes in vitro flowering in orchids (Kostenyuk et al., 199 9). Within the concentration ranges tested in the present study, treatments with 6-BA alone, 6-BA combined with either NAA or IBA, and the combined application of 6-BA, NAA, and IBA failed to induce floral bud formation in *C. tortisepalum* var. *longibracteatum*. This result is inconsistent with the findings of Hor et al. (2007) in *Dendrobium* ‘Chao Praya Smile’. TDZ is regarded as a unique plant growth regulator that exhibits both auxin-like and cytokinin-like activities, and its application can significantly alter endogenous auxin and cytokinin levels in plants (Guo et al., 2011). Ferreira et al. (2006) reported that TDZ promoted flower formation in *Dendrobium* species. In the present study, the combination of 0. 4 mg · L^−1^ TDZ + 1.0 mg · L^−1^ 6-BA + 0.8 mg · L^−1^ NAA + 1.6 mg · L^−1^ IBA produced the highest normal floral bud formation rate in *C. tortisepalum* var. *longibracteatum*, reaching 2 0.00%. The key finding is that TDZ was the key factor, with 0.4 mg · L^−1^ yielding the best results, and higher TDZ and 6-BA levels reduced bud formation. Range analysis of the normal floral bud formation rate identified TDZ as the primary contributing factor, followed by 6-BA. Compared with its closely related species, *C. goeringii*, the optimal TDZ concentration identified for C. *tortisepalum* var. *longibracteatum* in this study (0.4 mg · L^−1^) was similar to that reported for *C. goeringii* (0.3 mg · L^−1^) by Zhi et al. (2025). However, the optimal 6-BA concentration for C. *tortisepalum* var. *longibracteatum* (1.0 mg · L^−1^) was substantially lower than that reported for *C. goeringii* (9.0 mg · L^−1^), indicating differences in cytokinin sensitivity between these two closely related taxa. Furthermore, the normal floral bud formation rate obtained in the present study was slightly higher than the 16.00% reported for *C. goeringii*, suggesting that the hormone combination identified here is well suited for inducing in vitro flowering in *C. tortisepalum* var. *longibracteatum*.

### Effects of different concentrations of PP_333_ on in vitro flowering of C. tortisepalum var. longibracteatum

PP_333_, a gibberellin biosynthesis inhibitor, has been reported to effectively induce floral bud differentiation and promote flower development under in vitro conditions (Abdalla et al., 2021). In this study, we evaluated the effect of PP_333_ on in vitro flowering of *C. tortisepalum* var. *longibracteatum* by varying PP_333_ concentrations in a basal medium supplemented with 1.0 mg · L^−1^ 6-BA + 0.4 mg · L^−1^ TDZ + 0.8 mg · L^−1^ NAA + 1.6 m g · L^−1^ IBA. The results showed that floral bud induction decreased progressively with increasing PP_333_ concentration. However, all induced floral buds developed normally, resulting in relatively high rates of normal floral bud formation and a low incidence of abnormal flowers. These findings differ from those reported by Wang et al. (2009) in *Dendrobiumno bile*. However, they are consistent with the observations of Cen et al. (2010) in *Dendrobium officinale*, where 0.2 mg · L^−1^ PP_333_ yielded the highest flowering induction rate (90.90%), whereas increasing the concentration to 0.3 mg · L^−1^ inhibited flower induction. In *C. tortisepalum* var. *longibracteatum*, PP_333_ may suppress floral bud induction while reducing flower abnormalities because it lowers endogenous gibberellin levels. Because gibberellins are critical for floral organ elongation and development, moderate inhibition of gibberellin biosynthesis may limit flower stalk elongation and reduce floral abnormalities, whereas excessive inhibition may impede floral transition. Furthermore, these results suggest that different orchid species and genotypes may exhibit distinct sensitivities to PP_333_ and different interactions with cytokinins. Such differences may reflect endogenous hormone balance and signaling pathways, and the underlying molecular regulatory mechanisms warrant further investigation.

### Effects of GA_3_ pretreatment on in vitro flowering of C. tortisepalum var. longibracteatum

Research has shown that GA_3_ can effectively release plant dormancy and induce early flowering. Gedam et al.(2025) reported that 200 mg · L^−1^ GA_3_ significantly promoted early bolting and flowering in *Allium* tuberosum. Cardoso et al. (2012) found that 125 mg · L^−1^ GA_3_ improved both the normal floral bud rate and flower quality in *Phalaenopsis*. In the present study, plantlets pretreated with GA_3_ were subsequently transferred to a basal medium supplemented with 1.0 mg · L^−1^ 6-BA + 0.4 mg · L^−1^ TDZ + 0.8 mg · L^−1^ NAA + 1.6 mg · L^−1^ IBA. However, compared with those not pretreated with GA_3_, the flower bud induction rate was significantly lower, indicating a species-specific response that differs from that observed in *Allium* tuberosum (Gedam et al. 2025) and *Phalaenopsis* (Cardoso et al., 201 2). As a temperate terrestrial orchid, *C. tortisepalum* var. *longibracteatum* typically requires a low-temperature vernalization period to initiate flowering, so this discrepancy may reflect species-specific regulation of flowering. Exogenous GA_3_ can partially substitute for the cold signal, but its effective concentration window is narrow. The present study showed that 0.6 mg · L^−1^ GA_3_ effectively induced flower bud formation, whereas 1.0 mg · L^−1^ inhibited floral bud differentiation. This may be because high GA_3_ concentrations excessively promote vegetative growth, leaving insufficient photosynthetic products and hormonal signals for reproductive growth. These findings are consistent with those of Xue et al. (2024), who reported that higher GA_3_ concentrations inhibited floral bud differentiation in *Hydrangea paniculata* ‘Vanilla Strawberry’. Future studies may explore the combined application of GA_3_ with growth retardants such as PP333 or extend the low-temperature induction period after GA_3_ pretreatment to improve floral induction efficiency.

### Effects of ABA pretreatment on in vitro flowering of C. tortisepalum var. longibracteatum

ABA, an endogenous hormone that negatively regulates plant growth, is involved in processes such as protein and lipid synthesis, seed dehydration tolerance, seed dormancy, and flowering (Ting et al., 2015). In the present study, ABA pretreatment was followed by transfer to a basal medium supplemented with 1 mg · L^−1^ 6-BA + 0.4 mg · L^−1^ TDZ + 0.8 m g · L^−1^ NAA + 1.6 mg · L^−1^ IBA; however, only one floral bud was induced, indicating that ABA pretreatment had no significant effect on floral bud induction in this system. Consistent with the findings of Wang et al. (2002) in *Phalaenopsis*, ABA treatment significantly inhibited flower stalk emergence. Wang et al. (1997) reported that pre-culturing *Dendrobium officinale* on a medium containing ABA before transferring it to a 6-BA-containing medium significantly increased the flower bud induction rate. Similarly, Wang et al. (2006) found that pretreatment with PP_333_ and ABA increased the flower bud induction rate and the normal flowering rate of *Dendrobium moniliforme* to 93.3% and 80.0%, respectively. However, the results of the present study differed from those reported by *Dendrobium officinale* (Wang et al.,1997) and *Dendrobium moniliforme* (Wang et al., 2006) in that ABA pretreatment did not improve floral bud induction. It may be that differences in plant material account for variations in the response to exogenous ABA. In the present study, only a single ABA pretreatment was applied before transfer to a medium containing auxins and cytokinins. Different combinations of plant growth regulators may alter ABA’s regulatory effects. To date, the interactive effects between ABA and other plant growth regulators, as well as their optimal concentration ranges, have not been systematically evaluated. Therefore, the underlying regulatory mechanisms remain to be further elucidated.

## Conclusions

This study systematically elucidated the regulatory effects of plant growth regulators o n floral bud induction in in vitro plantlets of *C. tortisepalum* var. *longibracteatum* and established an efficient in vitro flowering system. TDZ was the key initiator of floral bud differentiation, as no floral buds were induced without it. Based on orthogonal experiments, the optimal hormone combination was 1.0 mg · L^−1^ 6-BA + 0.4 mg · L^−1^ TDZ + 0.8 mg · L^−1^ NAA + 1.6 mg · L^−1^ IBA, which resulted in a floral bud induction rate of 23.33% and a normal floral bud formation rate of 20.00%. PP_333_ inhibited floral bud differentiation, with induction rates decreasing as its concentration increased, while reducing the occurrence of abnormal flowers. GA_3_ pretreatment at 0.6 mg · L^−1^ for 60 days achieved a flower bud induction rate of 12.04%, whereas ABA pretreatment showed no significant promoting effect on flower bud induction. These results indicate that the optimized hormone combination and key parameters established in this study provide a practical technical framework and found ational data for shortening the breeding cycle of *C. tortisepalum* var. *longibracteatum* and for investigating the hormonal regulatory networks underlying floral transition in orchids.

## Acknowledgments

This work was funded by SiChuan Province Science and Technology Support Program (2017JY0132, 2024zd3302), LongShan Academic Talent Research Supporting Program (18LZX522).

## Abbreviations

1/2MS: half-strength Murashige and Skoog medium;
6-BA: 6-benzyladeni ne;
ABA: Abscisic acid;
GA_3_: Gibberellic acid;
IBA: Indole-3-butyric acid;
MS: Mura shige and Skoog medium;
NAA: α-naphthaleneacetic acid;
PP333: Paclobutrazol; TDZ - t hidiazuron.

