## Supplementary material for "Plant regulator of flower bud differentiation in *in vitro* plants of *Cymbidium tortisepalum* var. *longibracteatum* with TDZ as the key initiator": Table S1 Supplementary

Table S1 Suppl. Effects of ABA pretreatment on the in vitro flowering of *C. tortisepalum* var. *longibract eatum*. Mean ± SD, n = 3. Different lowercase letters within the same column indicate significant differ ences between treatments ($P$ < 0.05). CK - hormone-free medium.

| **Treatment** | **ABA(mg·L$^{-1}$)** | **Number of days of pre-treatment（d）** | **Flower bud induction rate（%）** | **Normal flower bud form ation rate（%）** |
|---|---|---|---|---|
| **CK** | Hormone-free MS medium | | 0.00a | 0.00a |
| **1** | 0.2 | 20 | 0.00a | 0.00a |
| **2** | 0.2 | 40 | 0.00a | 0.00a |
| **3** | 0.2 | 60 | 0.00a | 0.00a |
| **4** | 0.6 | 20 | 0.00a | 0.00a |
| **5** | 0.6 | 40 | 0.00a | 0.00a |
| **6** | 0.6 | 60 | 4.55±8.22a | 0.00a |
| **7** | 1.0 | 20 | 0.00a | 0.00a |
| **8** | 1.0 | 40 | 0.00a | 0.00a |
| **9** | 1.0 | 60 | 0.00a | 0.00a |
